# Accurate identification of cancer cells in complex pre-clinical models using deep-learning: a transfection-free approach

**DOI:** 10.1101/2023.10.15.562411

**Authors:** Marilisa Cortesi, Dongli Liu, Elyse Powell, Ellen Barlow, Kristina Warton, Caroline E. Ford

**Affiliations:** Gynaecological Cancer Research Group, School of Clinical Medicine, Faculty of Medicine and Health, University of New South Wales, Kensington, NSW, Australia; Laboratory of Cellular and Molecular Engineering, Department of Electrical Electronic and Information Engineering “G. Marconi”, Alma Mater Studiorum-University of Bologna, Cesena, Italy

## Abstract

3D co-cultures are key tools for *in vitro* biomedical research as they recapitulate more closely the *in vivo* environment, while allowing control of the density and type of cells included in the analysis, as well as the experimental conditions in which they are maintained. More widespread application of these models is hampered however by the limited technologies available for their analysis. The separation of the contribution of the different cell types, in particular, is a fundamental challenge.

In this work, we present ORACLE, a deep neural network trained to distinguish between ovarian cancer and healthy cells based on the shape of their nucleus. The extensive validation that we have conducted includes multiple cell lines and patient derived cultures to characterise the effect of all the major potential confounding factors. High accuracy and reliability were maintained throughout the analysis demonstrating ORACLE effectiveness with this detection and classification task.

ORACLE is freely available (https://github.com/MarilisaCortesi/ORACLE/tree/main) and can be used to recognise both ovarian cancer cell lines and primary patient-derived cells. This feature sets ORACLE apart from currently available analysis methods and opens the possibility of analysing *in vitro* co-cultures comprised solely of patient-derived cells.

## Introduction

3D cell culture is becoming a key tool of *in-vitro* research, enabling more accurate modelling of complex biological processes while maintaining a high level of control on the experimental system and its components (Jensen & Teng, 2020). The possibility of including multiple cell types within the same 3D culture is a fundamental advantage of these setups, as cell populations growing *in vivo* are not segregated and interact extensively with one another. The use of these co-cultures is however hampered by the limited techniques available for the identification of individual sub-populations.

Labelling of specific cell types by transfection with a fluorescent protein is currently the most common approach (Albrecht et al., 2006; Frimat et al., 2011; Gaté et al., 2022; Goers et al., 2014; Joshi et al., 2021; Peters et al., 2015a). While effective, this strategy has the notable drawback of restricting the analysis to commercial cell lines, which can be maintained in culture long enough to enable the screening and selection of fluorescently labelled cells. The production of fluorescent proteins has also been shown to be associated with a non-negligible metabolic burden which, in some cases, had measurable effects on cell behaviour (di Blasi et al., 2021). Recent advances in image analysis and machine learning could address this limitation and enable the label-free identification of different cell types. Indeed, multiple examples of software tools for the analysis of microscopy images are available and have been shown to be effective in improving the accuracy and reliability of the quantified variables (Cortesi et al., 2017, 2018; Lovecchio et al., 2019; Moen et al., 2019; Suarez-Arnedo et al., 2020).

This work focusses on ORACLE (OvaRian cAncer CeLl rEcognition), a freely available software tool for the classification of cancer and healthy cells from microscopy images. It is based on Faster-R-CNN, a deep neural network extensively used for object detection (Ren et al., 2015), which has shown accuracy and reliability in several domains (Benjdira et al., 2019; Singh et al., 2021; Su et al., 2021; Wan & Goudos, 2020). The target application presented in this work is the quantification of cell invasion in a 3D organotypic model of ovarian cancer metastasis. This model, initially described by Kenny and colleagues (Kenny et al., 2015) combines three different cell types (fibroblast, mesothelial and cancer cells) and mimics the structure of the omentum, the most common dissemination site for ovarian cancer. The three cell types are seeded in standard Transwell inserts with a surrogate extracellular matrix (Figure 1), enabling the quantification of cell invasion (Cortesi et al., 2023; Joshi et al., 2021; Peters et al., 2015b). Cancer cells used in these experiments, however, need to be transfected with a fluorescent protein to avoid overestimating invasion by including migrated healthy cells. ORACLE eliminates the need for this pre-emptive labelling of the cancer cells, being able to distinguish between healthy and cancer cells solely by the shape of the nucleus.

**Figure 1:**
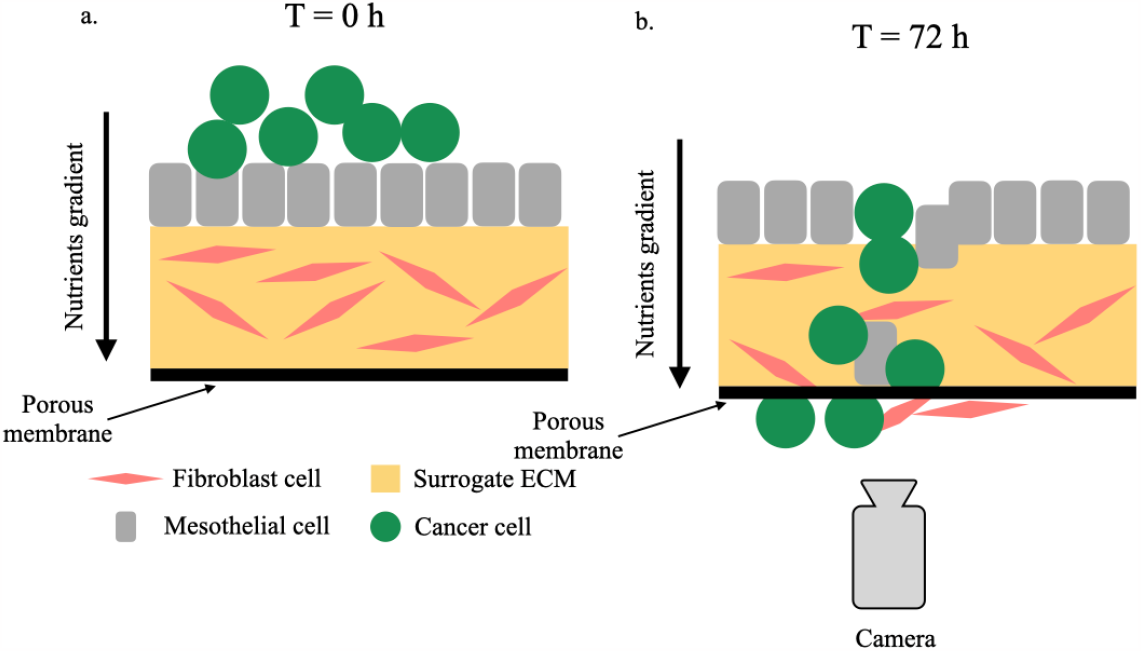
Schematic representation of the organotypic model and its use in Transwell invasion assays. a. Configuration at time 0. Cancer cells (green circles) are seeded on a layer of mesothelial cells (grey rectangles) which lays on a collagen surrogate extracellular matrix (ECM) containing fibroblasts. A nutrient gradient encourages cells to migrate downward. (b) At the end of the experiment images of the cells that have crossed the porous membrane, taken from the bottom of the culture, enable the quantification of cell invasion.

In this paper we describe the technical development of ORACLE, together with the extensive experimental validation that was conducted to verify its accuracy. The trained neural network, and the software used for training and data analysis are available at https://github.com/MarilisaCortesi/ORACLE/tree/main.

## Methods

### DNN construction and training

The pre-trained implementation of Faster R-CNN with a ResNet-50-FPN backbone available within the Pytorch library was used as a base for ORACLE (Ren et al., 2015). A total of 558 images (640x480 px, 55% cancer cells and 45% healthy cells), obtained by dividing the full-size images into 16 non-overlapping regions, were used to finetune the network’s training and enable the distinction of the two types of cells. The non-cancer class combined mesothelial and fibroblast cells, while the cancer class included images of two high grade serous ovarian cancer (HGSOC) cell lines (PEO1 and PEO4).

A data augmentation procedure was applied to increase the diversity of the dataset. Images were randomly flipped vertically and horizontally with a probability of 0.5. These transformations were chosen as they preserve cell size and shape proportions.

Training for 50 epochs (batch size 4) was conducted using an Adam optimiser with learning rate of 10^-4^. The coco-evaluator module was used to monitor performances on a test set comprising 62 images (55% cancer, 45% healthy).

Accuracy, precision and recall (Equations 1-3) were used to quantify ORACLE’s performance on the training and test set. Here TP, TN, FP and FN represent the true positive, true negatives, false positives and false negatives respectively.

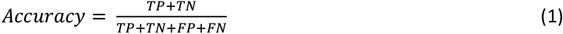

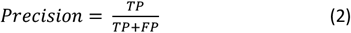

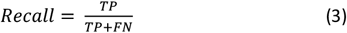

### Datasets and pre-processing

An extensive validation was then conducted, to evaluate the effect of each factor with the potential of influencing the results (Figure 2, Table 1).

**Table 1:** summary of the datasets used for ORACLE’s validation. The size of the dataset is the number of full-size images, prior to the division into sub-images.

| Dataset ID | Dataset name | Type of images | Size |
| --- | --- | --- | --- |
| 1 | No fluorescent tag | Immunohistochemistry | 16 images (291 cells) |
| 2 | Migration through the membrane | Fluorescence | 60 images (338 cells) |
| 3 | Different cell shapes | Immunohistochemistry | 52 images (2229 cells) |
| 4 | Co-culture with cell lines | Fluorescence | 222 images (10379 cells) |
| 5 | Primary cells | Fluorescence | 120 images (960 cells) |
| 6 | Co-culture with primary cells | Fluorescence | 50 images (7962 cells) |

**Figure 2:**
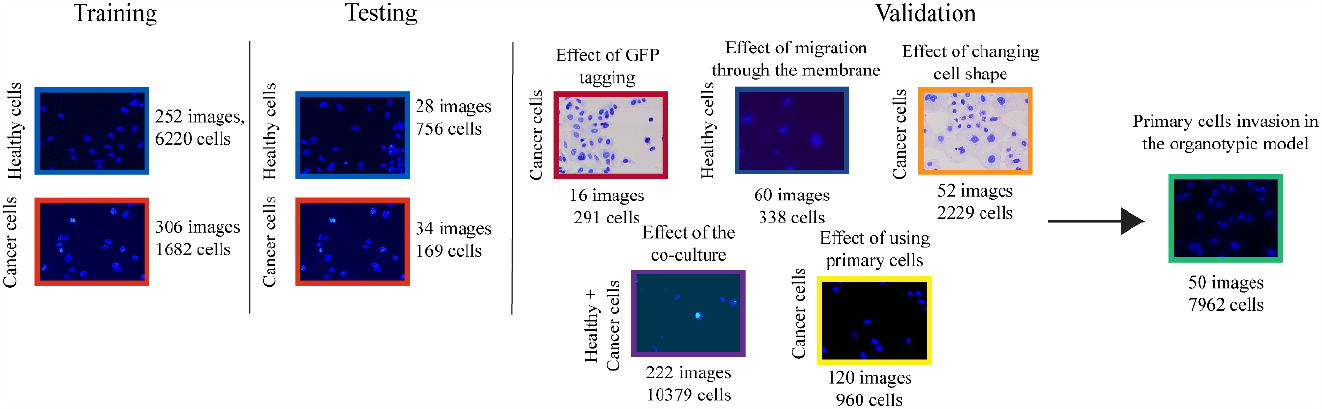
Summary of the analysis presented in this work. For each phase of ORACLE development and testing the size of the dataset and one representative image are reported.

To determine whether the addition of the fluorescent tag was altering the shape of the nucleus, we used hematoxylin-stained immunocytochemistry (ICC) images of the same HGSOC cell lines used during training (dataset ID 1). This method was also employed to evaluate the effect of changing cell shape, with 5 HGSOC cell lines (CaOV3, COV318, Kuramochi, OAW28, OVCAR4) that vary in cell morphology and size being tested (dataset ID 3). Images from Transwell assays conducted using a mixed population of fibroblast and mesothelial cells were used to evaluate the effect of migration through the membrane (dataset ID 2). Organotypic models using four different HGSOC cell lines tagged with GFP (CaOV3, OVCAR4, PEO1, PEO4) were then tested, to evaluate if co-culturing these cancer cell lines, and healthy cells would affect the results (dataset ID 4). Finally, Transwell assays using ascites-derived HGSOC cells were conducted (dataset ID 5), to check the system’s accuracy with the increased variability of patient-derived cells. Following these verification steps, a label-free organotypic model comprising of solely patient-derived cells (fibroblasts, mesothelial and ascites-derived cancer cells) was tested (dataset ID 6).

Given the differences between the datasets used in this work, customised pre-processing procedures were applied to each one, to improve their similarity to the training/test sets. For H&E images (datasets ID 1 and 3), the green channel of the brightfield images was inverted, and the exposure was adjusted though a gamma correction (gamma = 2 and gain = 1.5). The same exposure correction was applied to the fluorescence images (with the same parameters for datasets ID 2, 4, 5, and with gamma = 1 and gain =1.5 for dataset ID 6). For the healthy cells (dataset ID 2), the intensity values were also rescaled between the 2^nd^ and 98^th^ percentile of the contrast-adjusted image.

In all cases, the full-size images were split into 4 (immunohistochemistry images) or 16 (fluorescence images) regions of equal size, due to memory constraints. During the evaluation phase, low confidence boxes (<50%) were removed.

### Procedural segmentation

The annotations for the training and test datasets were obtained using the procedural segmentation previously described (Cortesi et al., 2023), which relies on a global thresholding using the Otsu’s method. The bounding boxes were then determined as the minimum and maximum values of the coordinates of each segmented cell. This software was also used to create the gold standard (i.e., the reference masks) for ORACLE’s validation (training, test and dataset ID 4). Manual screening of the images was conducted to exclude any incorrectly labelled ones and thus ensure the accuracy of the gold standard. As previously described (Cortesi et al., 2023) the segmented cells were classified as cancer if their area was between 50 and 5,000 pixels and they had an average fluorescence intensity higher than the background.

### Human research ethics approvals

Informed consent was obtained from all the patients involved in the study. The South Eastern Sydney Local Health District Human Research Ethics Committee (HREC) approved the protocol for the collection of the samples and their use in this project. Reference IDs for the collection of healthy and cancer (ascites derived) cells are SESLHD HREC approval #16/108 and Royal Hospital for Women HREC #19/001 respectively.

### Cell culture

CaOV3, PEO1, PEO4, and Kuramochi cells lines were kindly donated by Prof. Deborah Marsh, University Technology Sydney, and COV318 and OAW28 by Prof. Nikola Bowden, University of Newcastle. OVCAR4 cells were purchased from the American Type Culture Collection (USA). Cells were maintained in RPMI medium (Thermo Fisher, Waltham, MA, USA), supplemented with 10% FBS (Scientifix, Australia), 1% Pen-strep (Sigma-Aldrich, USA) and 1% GlutaMAX (Thermo Fisher, Waltham, MA, USA). All cell lines were confirmed free of mycoplasma infection and validated via STR profiling. When required, transfection with the pLKO.1-Neo-CMV-tGFP vector (Sigma-Aldrich, USA) was used to label cells with GFP.

Fibroblasts and mesothelial cells were extracted from omentum samples collected from patients undergoing surgery for benign or non-metastatic conditions at the Royal Hospital for Women and Prince of Wales Private Hospital (site specific approval ethics # LNR/16/POWH/236). The tissue was processed as previously described (Kenny et al., 2015) and the isolated cell populations were maintained in DMEM media supplemented with 10% FBS (Sigma-Aldrich, USA), 1% non-essential amino acids (Sigma-Aldrich, USA) and 1% GlutaMAX (Thermo Fisher, Waltham, MA, USA) and 1% Antibiotic-Antimycotic (Gibco, USA).

Ascites samples were collected from patients undergoing paracentesis at the Royal Hospital for Women following a HGSOC diagnosis. Cancer cell cultures were obtained by adding 25 ml of ascites fluid to a T175 cell culture flask and progressively substituting it with cell culture media over the course of a few days. Adherent cells were then maintained in RPMI medium (Thermo Fisher, Waltham, MA, USA), supplemented with 10% FBS (Sigma-Aldrich, USA), 1% Pen-strep (Sigma-Aldrich, USA) and 1% GlutaMAX (Thermo Fisher, Waltham, MA, USA). The study applied initial passages (up to passage 3 to avoid inducing morphological alterations).

### Immunocytochemistry and immunofluorescence

HGSOC cell lines (CaOV3, COV318, Kuramochi, OAW28, OVCAR4) were seeded in 8-well chamber slides (Thermo Scientific™ Nunc™ Lab-Tek™ II Chamber Slide™) at a 70% confluency. After overnight adhesion, cells were fixed with 4% paraformaldehyde, permeabilised with 0.5% Triton X100, and counterstained with hematoxylin. The slides were mounted with coverslips and visualised under the light microscope.

Fibroblasts and mesothelial cells from 2 donors, were seeded separately at a 70% confluency in 8-wells chamber slides (Thermo Scientific™ Nunc™ Lab-Tek™ II Chamber Slide™). After 24h of incubation, cells were fixed with 4% paraformaldehyde and the slide was mounted using a medium containing 4′,6-diamidino-2-phenylindole-DAPI (Fluoroshield, Sigma-Aldrich, USA). A fluorescence microscopy setup (Leica DM 2000 LED fitted with a Leica DFC450c camera) was used to acquire the images. A total of 6 wells (3 per cell type) were acquired.

### Invasion assay

Transwell invasion assays were conducted as previously described (Cortesi et al., 2023). Briefly, for monoculture models (dataset ID 2, 5 and training and test sets) Matrigel-precoated Transwell inserts (Corning Life Sciences, US) were seeded with either HGSOC cells (Table 2) or a mixture of fibroblasts and mesothelial cells (Table 2). Samples were incubated for 24 or 48 h (Table 2) during which a chemoattractant (FBS) was used to induce cell invasion and migration through the membrane.

**Table 2:** Characteristics of the Transwell invasion assays conducted in this work. Experimental conditions (seeding density and incubation time) were optimised independently for each dataset to maintain the number of invaded cells/image within a reasonable range.

| Dataset | Cells | Seeding density | Incubation time |
| --- | --- | --- | --- |
| Training/test set | PEO1 | 2M cells/ml | 24 h |
| Training/test set | PEO4 | 1M cells/ml | 48 h |
| 2 | Fibroblasts and mesothelial cells | 40K cells/ml (fibroblasts),<br>200K cells/ml (mesothelial) | 48 h |
| 4 | Co-culture of HGSOC cell lines and healthy cells | 40K cells/ml (fibroblasts),<br>200K cells/ml (mesothelial)<br>2M cells/ml (cancer) | 24 h (PEO1), 48 h (PEO4, CaOV3, OVCAR4) |
| 5 | Ascites-derived HGSOC cells | 1M cells/ml, 2M cells/ml | 24 h |
| 6 | Co-culture of ascites derived HGSOC cells and healthy cells | 40K cell/ml (fibroblasts),<br>200K cells/ml (mesothelial)<br>2M cells/ml (cancer) | 24 h |

For the co-culture models, regular Transwell chambers were used (Corning Life Sciences, US). A layer of fibroblasts (4·10^4^ cells/ml) embedded in a solution of collagen (5 ng/μl, Sigma-Aldrich, USA, in 100 μl of culture media) was seeded at the bottom of the Transwell insert. After 4 h of incubation in standard culturing conditions (37ºC and 5% CO_2_), 50 μl of media containing 20,000 mesothelial cells was added on top and incubated from 24 hours to allow a monolayer of mesothelial cells to form. Cancer cells were seeded on the mesothelial layer after 24 h (Hart et al., 2021; Kenny et al., 2015; Peters et al., 2015b), at which point a nutrient gradient was created by using medium with a low FBS concentration (1%) in the in the insert, while the bottom chamber was filled with 20% FBS medium. Incubation times for the co-culture datasets are reported in Table 2.

At the end of the experiment, Transwell inserts were fixed with 100% methanol and the membranes were mounted on microscope slides using a medium containing DAPI (Fluoroshield, Sigma-Aldrich, USA). 10 images from different regions of the membrane were acquired with a fluorescence microscopy setup (Leica DM 2000 LED fitted with a Leica DFC450c camera).

### Statistical analysis

The non-parametric Kolmogorov-Smirnov test was used, whenever appropriate, to compare different conditions. A p-value of 0.05 was considered as threshold for significance.

## Results

### Training of the network

ORACLE was built by finetuning a pre-trained Faster R-CNN network. This solution was preferred to de-novo training, as it requires less data to achieve good detection and classification performances (Weiss et al., 2016). Indeed, less than 600 images from two HGSOC cell lines (PEO1, PEO4) and healthy (mesothelial and fibroblast) cells were sufficient to achieve effective detection and classification of the two cell types (Table 3). Equivalent results were obtained when analysing the test set (Table 3), which consisted of 62 images of the same cell type acquired with the same techniques as the training set, that is, Transwell assays for the cancer cells and immunofluorescent staining for fibroblasts and mesothelial cells.

**Table 3:** Performance metrics calculated for the training and test set. The definitions of accuracy precision and recall are reported in the methods.

|  | Training Set |  | Test Set |  |
| --- | --- | --- | --- | --- |
|  | Healthy cells | Cancer cells | Healthy cells | Cancer cells |
| Accuracy | 0.95 +/- 0.12 | 0.92 +/- 0.17 | 0.97 +/- 0.08 | 0.95 +/- 0.09 |
| Precision | 0.95 +/- 0.19 | 0.93 +/- 0.23 | 1.0 +/- 0.02 | 0.97 +/- 0.07 |
| Recall | 0.91 +/- 0.15 | 0.89 +/- 0.18 | 0.94 +/- 0.10 | 0.95 +/- 0.09 |

Figure 3a shows an example of the results obtained with ORACLE. Each region of the image recognised as a cell is outlined by a bounding box, colour-coded according to the corresponding classification, red for cancer and blue for healthy cells. Figure 3b shows the comparison between ORACLE’s results and the gold standard values obtained using the procedural segmentation described in (Cortesi et al., 2023). This algorithm relies on morphological and intensity features to distinguish between cancer and healthy cells.

**Figure 3:**
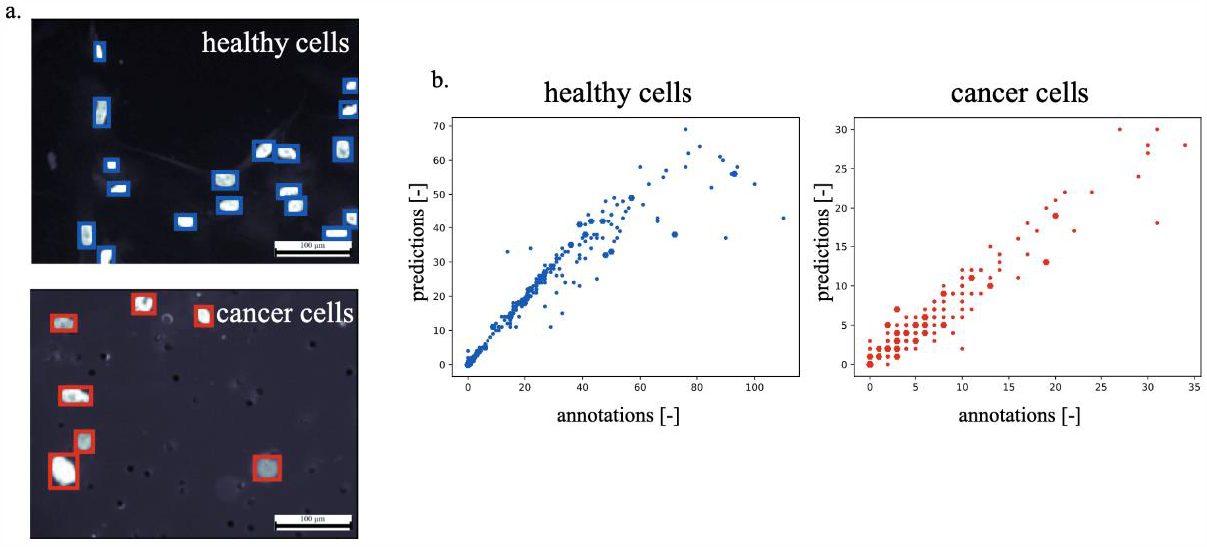
Results of ORACLE training. a. Representative images of the healthy and cancer cells used for the training and testing of the DNN (magnification 20x). The overlayed boxes outline the predicted cells and are colour coded according to the predicted class (red for cancer and blue for healthy). b. Comparison of the annotated and predicted number of cells for each class. Blue refers to the healthy cells (R^2^ = 0.96) while red refers to the cancer cells (R^2^ = 0.97). Dots identify the images of the training set, while hexagons the ones of the test set.

The Pearson’s correlation coefficient was very high (above 0.95 and p<0.01 for both classes), and there was no noticeable difference between training (dots) and testing (hexagons) sets. This analysis, beside confirming the results in Table 3 at the image level, demonstrates the effectiveness of ORACLE in distinguishing between healthy and cancer cells in images of monocultures of these cell types.

### Effect of the fluorescent tag of cancer cells recognition

To verify that the nucleus shape was not affected by the addition of the fluorescent tag, ORACLE was tested on a set of images of wild-type PEO1 and PEO4 cells acquired using a brightfield microscope following hematoxylin staining (left side of Figure 4a). These images were pre-elaborated as described in the methods to match the features of the training/test sets (clear, bright cells and dark background). The results of this procedure are reported in Figure 4a, where the same image is shown both in its original (left side) and pre-elaborated (right side) version. In this figure, the bounding boxes outline each of the recognised cells. As for Figure 3, red identifies cancer cells while blue identifies the healthy cells.

**Figure 4:**
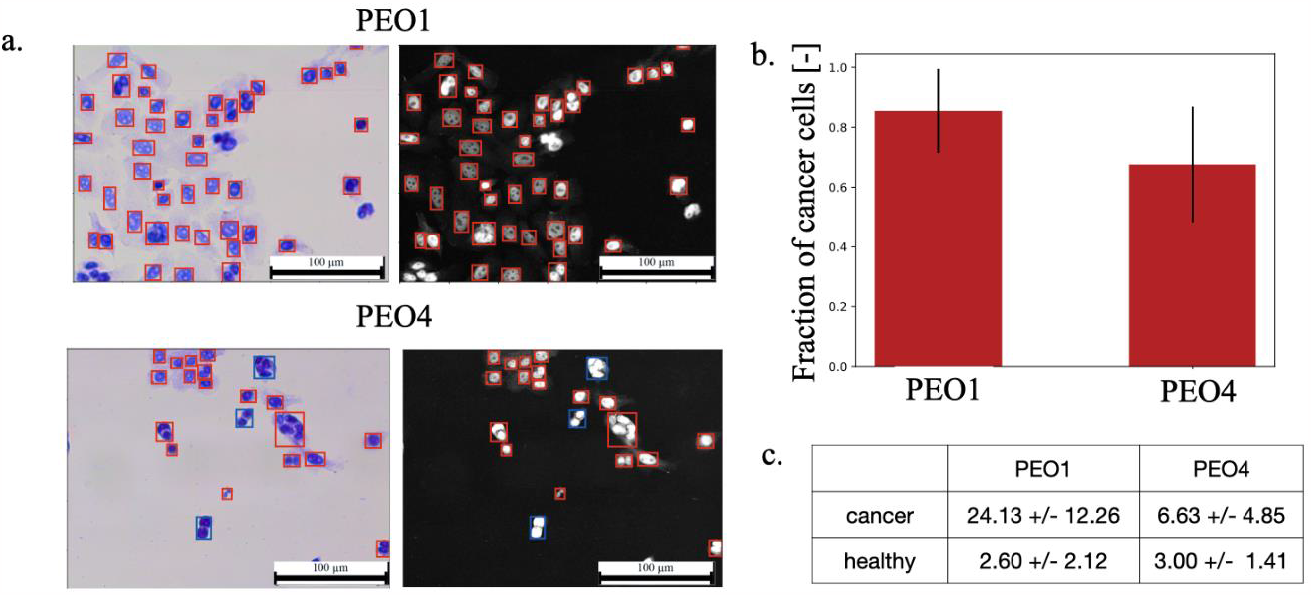
a. Representative images of PEO1 and PEO4 cells without GFP tag (magnification 20x). On the left side there is the original image, while the right hand shows the result of the pre-elaboration. Overlayed boxes outline the predicted cells and are colour coded according to the predicted class (red for cancer and blue for healthy). b. Fraction of cells correctly classified as cancer for each cell line. Data represented as mean +/- standard deviation. c. Table showing the average (+/- standard deviation) of the number of cells of each class for each cell line.

The fraction of cancer cells in Figure 4b provides a measure of the accuracy of the classification, as no healthy cells were used for this experiment. Most cells were classified correctly, but the fraction of cancer cells was, on average, 18% higher for PEO1. This is unlikely to be indicative of a difference in detection accuracy, as both PEO1 and PEO4 were used as examples for cancer cells during training and were equally represented in the dataset. Additionally, images containing PEO4 contained fewer cells (Figure 4c) while the number of misclassified cells was on average 3 per image for both cell lines. This inevitably reduced the fraction of cells correctly classified for the PEO4 condition as the same number of mis-classified objects represents a higher fraction of the total.

These results confirm the effectiveness of ORACLE in identifying cancer cells which were not fluorescently tagged. They also suggest that using a medium/high cell density might be advisable, as the rate of error shows a weak dependence on the number of cells in the image.

### Effect of migration through the membrane on nucleus shape of non-cancer cells

To investigate whether migration through the membrane affects the shape of the nucleus and thus ORACLE’s accuracy, we conducted Transwell migration assays using only healthy cells, in the same density and proportion as in the organotypic model. Samples from two different patients were used in this analysis, to account for individual variability.

Figure 5a shows representative images from each of the patients with bounding boxes outlining the recognised cells. Colours are used to identify the corresponding class (in this case only blue for healthy cells). The fraction of correctly classified cells was used as a metric of accuracy (Figure 5b), together with the overall number of cells for each class (Figure 5c).

**Figure 5:**
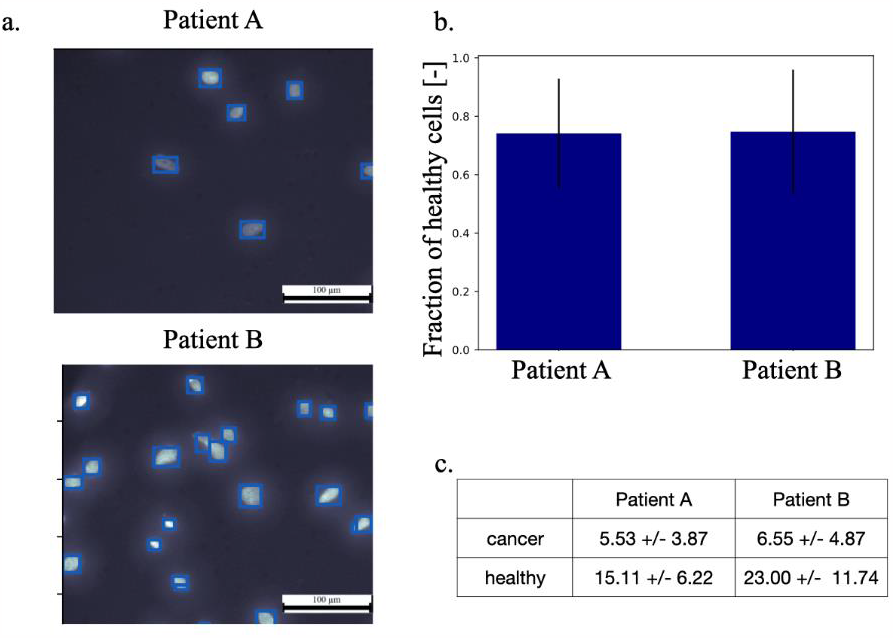
a. Representative images of fibroblasts and mesothelial cells that have invaded through the membrane (magnification 20x). Overlayed boxes outline the predicted cells and are colour coded according to the predicted class (blue: healthy). b. Fraction of cells correctly classified as healthy for each patient. Data represented as mean +/- standard deviation. c. Table showing the average (+/- standard deviation) of cell numbers for each class and patient.

Nucleus morphology was shown to be largely conserved between patients, with no discernible difference in the raw images or in the classification accuracy (Figures 5a and 5b). The cells derived from patient B, however, were characterised by a higher invasion with on average 10 cells/image more than what measured for patient A. This difference was mainly determined by the variation in the number of cells classified as healthy, as the rate of misclassification was approximately constant (Figure 5c).

The high classification accuracy (about 75%), and its negligible dependence on patient specific features, demonstrate the effectiveness of ORACLE in recognizing healthy cells even after they have crossed the membrane of the transwell insert.

### Effect of changing the cancer cells lines

Generalisation beyond PEO1 and PEO4 cells is another requirement for ORACLE as several HGSOC cell lines are available and currently used in research. To verify this aspect hematoxylin stained images of 5 different HGSOC cell lines of variable shape and size (CaOV3, COV318, Kuramochi, OAW28, OVCAR4) were considered and the fraction of cells correctly classified as cancer was used as metric. The brightfield images composing this dataset were pre-elaborated, using the same procedure as the ones in Figure 4, to maximise their similarity to the training/test set (Figure 6).

**Figure 6.**
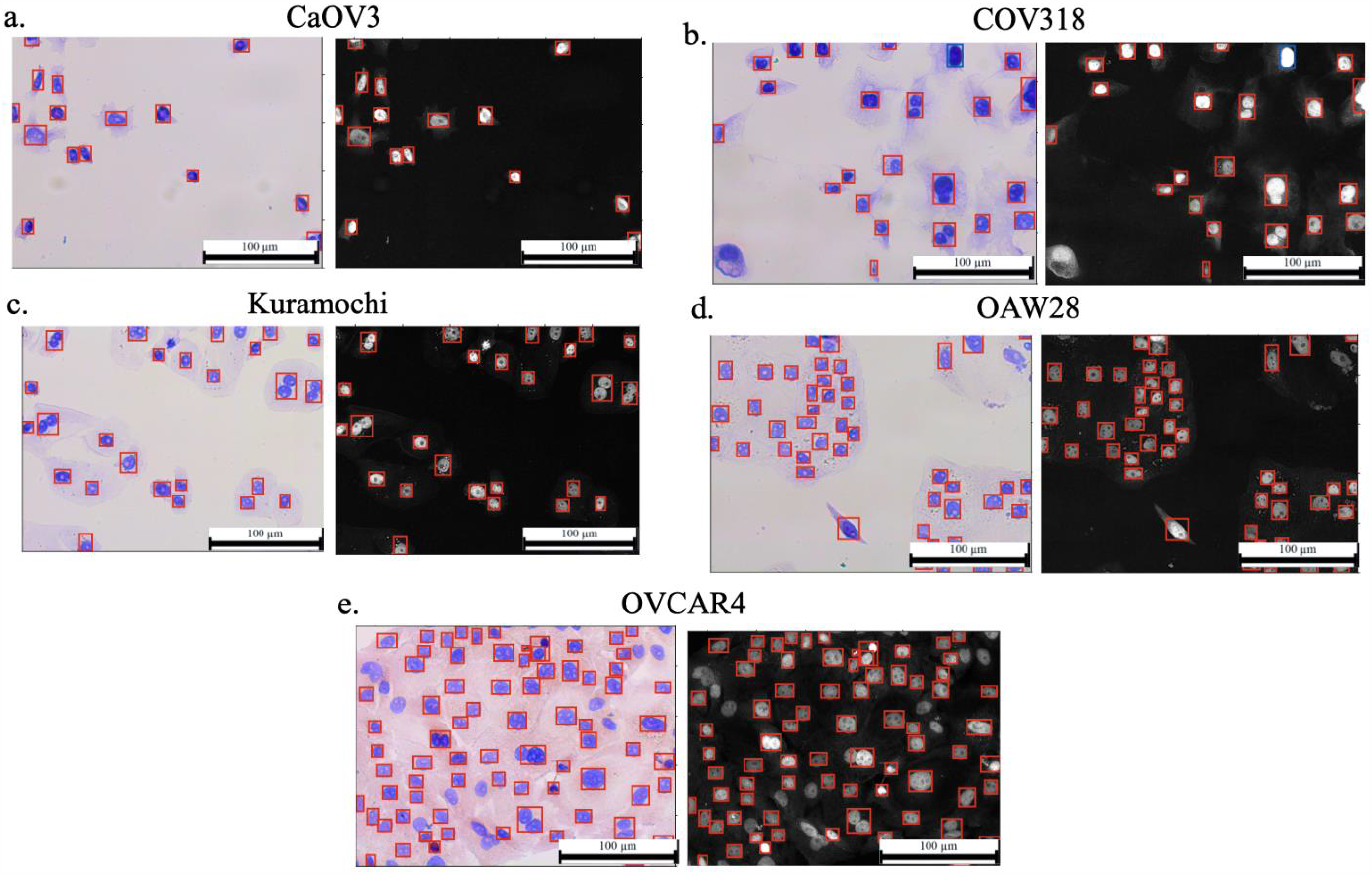
a.- e. Representative images of the five other cell lines used in this work (Magnification 20x). On the left-hand side there is the original image, while the right-hand side shows the result of the pre-elaboration. The overlayed boxes outline the predicted cells and are colour coded according to the predicted class (red: cancer, blue: healthy).

The fraction of correctly classified cells (Figure 7a), together with the distributions of the number of cells for each class (Figure 7b) confirm the ability of ORACLE of generalising beyond PEO1 and PEO4. Indeed, a very high accuracy (above 80%) was retrieved in most cases, with at least a 5-fold difference between the number of correctly recognised and misclassified cells. Images of OVCAR4 cell were the only ones associated with a reduced accuracy (below 60% on average) and a larger proportion of incorrectly classified cells (Figure 7b). The higher cell density observed for images of OVCAR4 cells, together with its high level of spatial organization and larger cytoplasm (Figure 6e) were the main determinants of this result, as further detailed in the next section.

**Figure 7:**
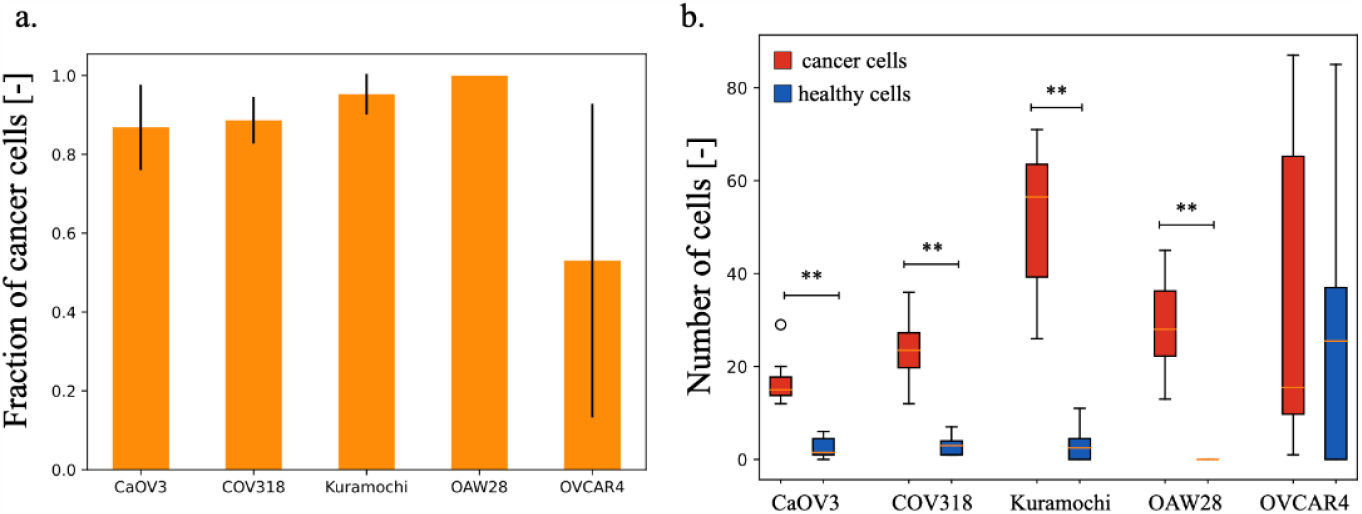
a. Fraction of cells correctly classified as cancer for each cell line. Data represented as mean +/- standard deviation. b. Boxplots showing the distribution of cell numbers for each class and cell line (Kolmogorov Smirnov test **: p < 0.005, * p< 0.05).

### Effect of co-culturing healthy and cancer cells

The organotypic model described by Kenny and colleagues (Kenny et al., 2015) was used for this step, in combination with 4 different HGSOC cell lines: PEO1, PEO4, CaOV3 and OVCAR4. The first two were chosen as they were used during the training of the network, while CaOV3 and OVCAR4 were included as examples of cells, respectively, easy and difficult to classify using ORACLE. In all cases, cancer cells were genetically modified to produce GFP, to enable their distinction from fibroblast and mesothelial cells so that reference cell counts could be obtained. The gold standard cell counts were obtained using the procedural segmentation algorithm presented by Cortesi and colleagues (Cortesi et al., 2023).

Figure 8 shows the results of this analysis. The number of cells per image varied widely, as shown in the representative images in panel a. This did not affect the accuracy of the analysis. Indeed, the average number of cells measured using ORACLE was highly consistent with the results of the procedural segmentation (Figure 8b). Most of the cells that invaded through the membrane were classified as cancer cells (Figure 8c). This is consistent with the increased aggressiveness often measured when maintaining cancer cells in 3D environments (Azimi et al., 2020; Cortesi et al., 2023; Pasini et al., 2021). In this dataset, OVCAR4 cells were not characterised by a reduced detection accuracy. This suggests that the results in Figure 7 were likely affected by the cytoplasm staining and the high cell density. Overall, these results confirm ORACLES’s ability to differentiate between the nuclei of cancer and healthy cells in co-culture conditions.

**Figure 8:**
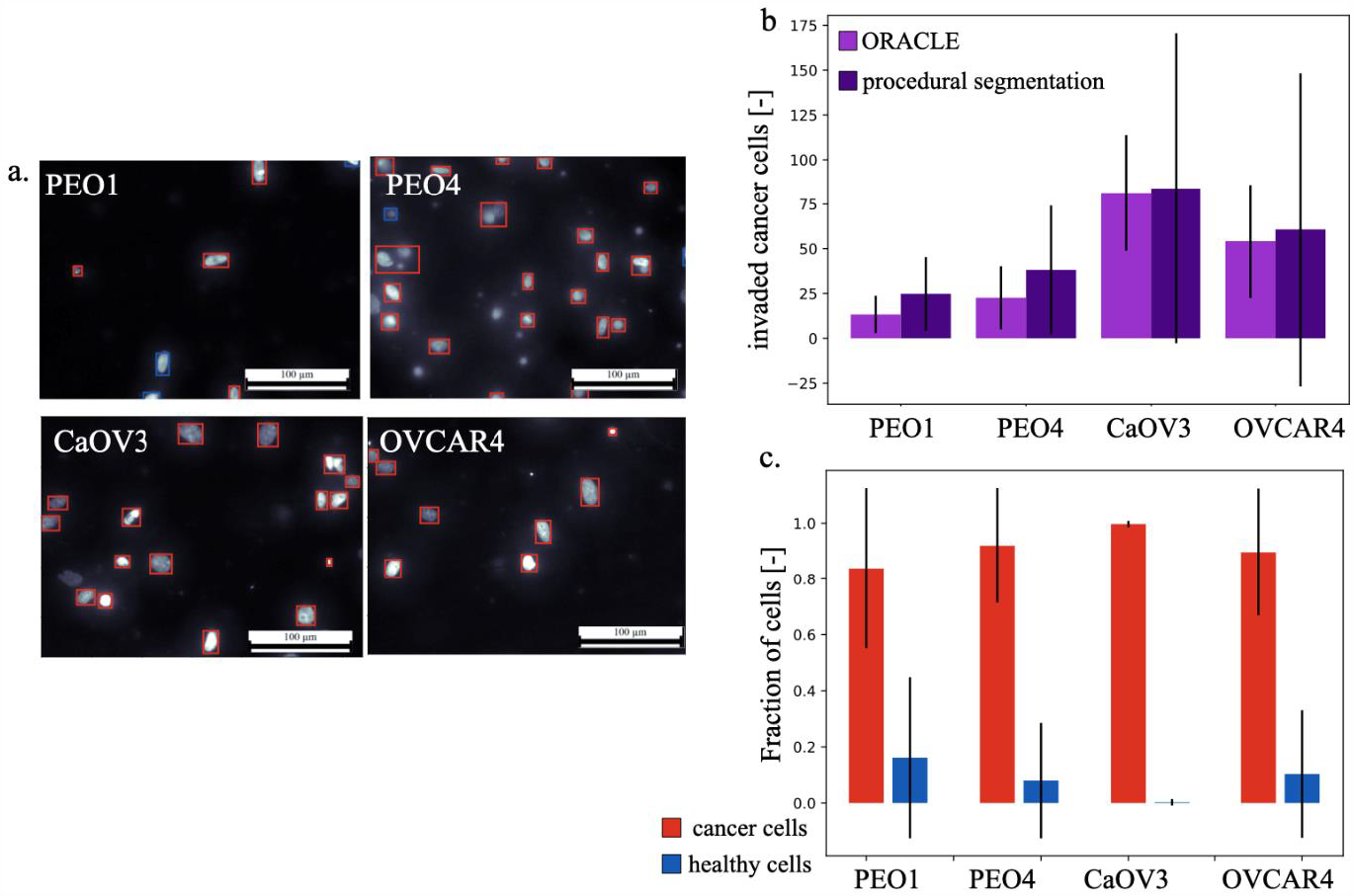
a. Representative image of the dataset analysed in this section (Magnification 20x). The overlayed boxes outline the predicted cells and are color coded according to the predicted class (red: cancer, blue: healthy). b. Average (+/- standard deviation) of the number of cancer cells obtained, for each condition with ORACLE (light purple) and the procedural segmentation algorithm used as reference (dark purple). c. fraction of cells classified by ORACLE as either healthy or cancerous for each cell line (data represented as mean +/- standard deviation).

### Effect of using patient-derived cancer cells

Transwell assays using a monoculture of ascites derived HGSOC cells were conducted. Samples from two different patients were considered, to evaluate whether ORACLE’s performance was dependent on patient-specific cell features. We also tested two different seeding densities (1M and 2M cells/ml) to evaluate the dependence of the results on the number of cells per image.

Figure 9 reports the results of this analysis. In panel a, one reference image for each condition is shown, with overlayed bounding boxes outlining the recognised cells. Red boxes identify the cells classified as cancer while blue is used for healthy cells. Almost all cells were correctly identified as cancer (Figure 9b), despite the significant variability observed between the tested conditions (Figure 9c). Indeed, the range of cancer cells/image varies over 3-fold and a differential dependence on the seeding density was observed for each patient. Cells derived from patient D invaded through the membrane at a rate that was almost independent from the seeding density, as only a 16% increase in the number of cells/image was observed doubling the seeding density. The same change in initial population cardinality resulted, on the other hand, in an approximate doubling of the average number of invaded cells for the samples obtained from Patient C. This is consistent with a constant invasion rate.

**Figure 9:**
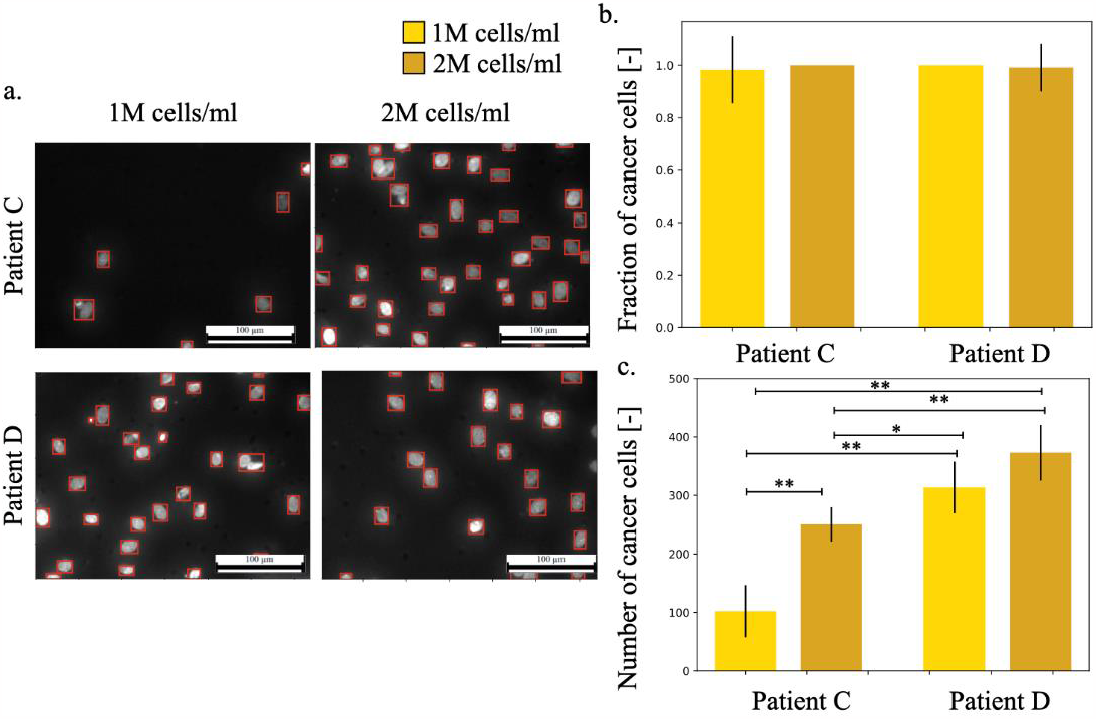
a. Representative images (magnification 20x) for all tested conditions (two patient derived HGSOC cancer cells seeded at two different densities). The overlayed boxes outline the predicted cells and are colour coded according to the predicted class (red: cancer, blue: healthy). b. Fraction of cells correctly classified as cancer for each condition. Data represented as mean +/- standard deviation. c. Average number of cancer cells/image obtained for each condition (Kolmogorov Smirnov test *: p < 0.05, **: p < 0.005).

While further investigation would be needed to determine the cause of this difference and its potential role in patient outcome and disease progression, this analysis demonstrates that ORACLE’s accuracy is maintained when using patient derived HGSOC cells.

### Study of the invasion of patient-derived cells in the organotypic model

Finally, we conducted an experiment in which two different patient-derived HGSOC cancer cell preparations were seeded on the organotypic model. To the best of the authors’ knowledge, this is the first time this analysis has been conducted, due to the challenges of distinguishing different cell types through the standard quantification of Transwell assays.

We tested both cells that had been previously used in ORACLE’s validation (combination 1: fibroblasts from patient A, mesothelial cells from patient B and cancer cells from patient D) and entirely novel samples (combination 2) to further test ORACLE’s generalisation abilities.

Figure 10 shows the results of this analysis. Most cells were classified as cancer (Figure 10b), coherently with the results obtained when cell lines were used (Fig 8 c). The absolute number of cancer cells was also consistent across the two considered samples, even though combination 1 had a higher variability. These results demonstrate the feasibility of using a DNN to identify different cell types in images of co-cultured cells. This greatly expands the scope of the co-culture model of HGSOC used in this study and opens to the possibility of developing similar tools for other complex culture systems.

**Figure 10:**
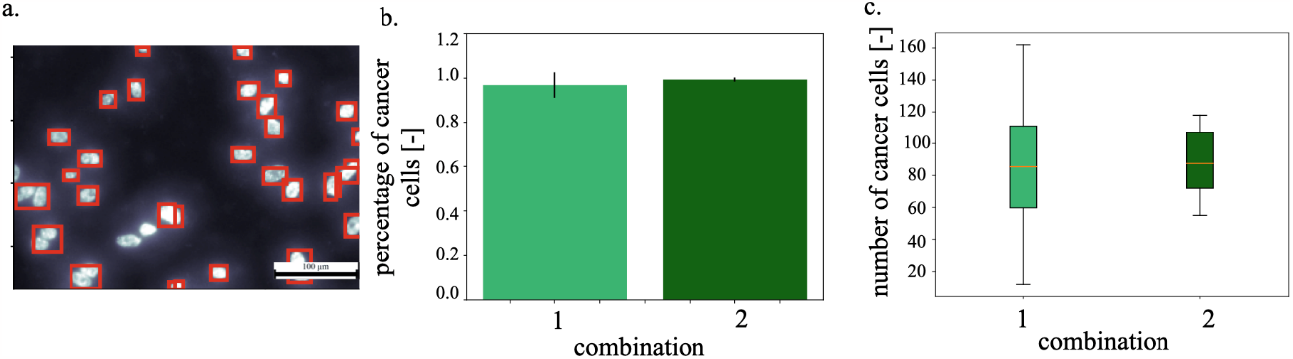
a. Representative image for the co-culture experiment comprising solely patient-derived cells (magnification 20x). Overlayed boxes outline the predicted cells and are colour coded according to the predicted class (red: cancer, blue: healthy). b Fraction of cells classified as cancer for each combination of cells. Data represented as mean +/- standard deviation. c. Distributions of the number of cancer cells/image obtained for each combination of cells.

## Discussion

3D co-cultures represent a key step in the development of more accurate *in vitro* models, as they enable study of the interaction among different cell populations and between cells and their environment. Separating the contribution of the different cell types, however, has been a challenge as most assays developed for monoculture models lack the ability to generalise to co-culture settings. Inducing the production, in specific cell types, of fluorescent markers is a solution, but it increases the complexity of the assay, and it requires the continuous selection of the culture with antibiotics. This stress, together with the burden associated with the production of the fluorescent and antibiotic resistance proteins, has been shown to affect cell behaviour in a largely unpredictable manner (di Blasi et al., 2021). The time required for the establishment of a stable transfected culture, additionally, is generally not compatible with the use of patient-derived cells, thus restricting the analysis to traditional cell lines.

In this work, we have presented ORACLE, a DNN trained to distinguish between HGSOC cells and healthy omentum cells (fibroblast and mesothelial cells) solely by the shape of the nucleus. Staining with DAPI was used to visualise the nucleus, as it is fast, widely available, and has a high signal to noise ratio. Another advantage of DAPI is that it can be used reliably on any cell, and is applied only at the end of the experiment, thus avoiding any potential influence on cell behaviour.

ORACLE is also an automatic quantification method. As such its results do not depend on the operator conducting the analysis and their ability to accurately identify and classify cells. This feature addresses one of the main limitations of manual counting, the most common analysis approach for Transwell assays, and improves the reliability and accuracy of the analysis. A further advantage of automation is the reduced workload of increasing the number of images. Indeed, while the elaboration time will be proportional to the size of the dataset, the limited operator involvement opens to the possibility of further strengthening the results through the acquisition of more images.

The extensive validation described in this work was developed to characterise the effect of all the major factors potentially influencing ORACLE’s performance. We tested cancer cells that had not been tagged with GFP, to evaluate whether transfection would affect the shape of the nucleus.

HGSOC cell lines not included in the training and patient-derived cultures were used to study the robustness of the DNN to changes in cell shape and intra-culture variability. Images of healthy cells that had migrated through the membrane were also considered, as the process of invasion is often associated with morphological changes facilitating cell motility and could affect classification accuracy. Co-cultures of healthy cells and cancer cells tagged with GFP were also included, to measure the performance of ORACLE on mixed populations. A very good accuracy was maintained throughout the analysis demonstrating the feasibility of using this tool for the quantification of HGSOC cell invasion in a 3D co-culture setting.

This result opens the possibility of analysing co-cultures comprised solely of patient-derived cells, an application currently not supported by any other method. This innovation holds great potential, as maintaining cells in a 3D co-culture induces measurable changes in their ability to invade. Indeed, cancer cells extracted from patient D showed a more pronounced invasion when cultured alone (Figure 9 c) when compared to the results obtained in the co-culture model (combination 1 in Figure 10 c). As the percentage of cells classified as cancer was consistent between the two experiments (Figure 9b and 10b), we can exclude misclassification as the source of this difference. While further analysis would be required to characterise this phenomenon, the preliminary study presented in this work highlights the limitations of the results obtained using cell lines. Indeed, PEO4 cells were previously shown to exhibit the opposite behaviour, that is higher invasion when maintained in a co-culture with fibroblasts and mesothelial cells (Cortesi et al., 2023).

The patient-specific variability observed in this analysis further highlights the need for more extensive use of patient-derived cell populations in preclinical research. Indeed, each cell population was characterised by a different invasion pattern. Patient C was associated with a constant invasion rate, which led to a direct proportionality between the cancer cell seeding density and the number of invaded cells. Patient D, on the other hand, exhibited a limited dependence on the initial condition and a reduction in invasion when cells were maintained in the organotypic model. Co-culturing with fibroblasts and mesothelial cells was also associated with an increase in variability.

Indeed, the ratio between the standard deviation and the mean is on average 2-fold higher for the data in Figure10, when compared with Figure 9. This effect was more pronounced for combination 1, which included the cancer cells extracted from patient D. While no indication of the *in vivo* behaviour of these cells is available, the differences between co-culture and monoculture results further highlights the importance of the experimental setting for the *in vitro* study of invasion.

A more extensive evaluation of the behaviour of healthy cells at the metastasis site is also warranted. Indeed, the two sets of fibroblast and mesothelial cells used for the analysis in Figure 5 yielded statistically different results. Further analysis will be required to fully characterise this difference and its effect on cancer cells behaviour, but this work provides the framework for this study and the analysis tool required to separate the contribution of healthy and cancer cells.

Finally, ORACLE opens the possibility of analysing invasion of HGSOC cells in surrogate tissues comprising multiple cell types derived from the same patient. This would enable the *in vitro* study of the role of cell-cell interaction in HGSOC metastasis in a setting closely resembling *in vivo* condition. At the same time, the use of an organotypic model allows control of the composition of the culture thus making it possible to characterise the role of each cell type in invasion and metastasis, together with the effect of changing the surrogate ECM.

This approach is not limited to organotypic models either. As shown in Figures 4 and 6, it can be extended to immunocytochemistry images, after an appropriate pre-elaboration. Use in other complex cell culture models, including organoids, tumoroids and explants, is also possible, provided that the images obtained from these cultures maintain a good similarity with the training dataset. Re-training with more representative examples is also possible and mainly limited by data availability. This flexibility is a major advantage of ORACLE that is expected to increase and expand its impact and usefulness for the scientific community.

## Conclusion

DNNs have revolutionised image analysis, greatly improving the reliability and accuracy of object detection and classification tasks. Harnessing this innovation in biomedical research holds significant great potential, both in terms of improving the analysis of already available assays and developing novel techniques. ORACLE is a good example in this regard, as it addresses the main limitations of the analysis of Transwell assays while expanding its scope of application. Indeed, it enables the study of invasion in co-cultures comprising solely of patient-derived cells, supports an increase in throughput for this technique and the characterisation of individual variability in HGSOC invasion and metastasis. At the same time, it relies on widely available instrumentation and common reagents. This substantially increases ORACLE’s base of potential users and supports the development of similar tools for other applications.

